# Stabilization of β-catenin promotes melanocyte specification at the expense of the Schwann cell lineage

**DOI:** 10.1101/2020.06.29.179291

**Authors:** Sophie Colombo, Valérie Petit, Roselyne Y Wagner, Delphine Champeval, Ichiro Yajima, Franck Gesbert, Irwin Davidson, Véronique Delmas, Lionel Larue

## Abstract

The canonical Wnt/β-catenin pathway governs a multitude of developmental processes in various cell lineages, including the melanocyte lineage. Indeed, β-catenin regulates *Mitf-M* transcription, the master regulator of this lineage. The first wave of melanocytes to colonize the skin is directly derived from neural crest cells, while a small number of second wave melanocytes is derived from Schwann-cell precursors (SCPs). We investigated the influence of β-catenin in the development of melanocytes of the first and second waves by generating mice expressing a constitutively active form of β-catenin in cells expressing tyrosinase. Constitutive activation of β-catenin did not affect the development of truncal melanoblasts, but led to a marked hyperpigmentation of the paws. By activating β-catenin at various stages of development (E8.5-E11.5), we showed that the activation of β-catenin in bipotent SCPs favored melanoblast specification at the expense of Schwann cells in the limbs within a specific temporal window. In addition, hyperactivation of the Wnt/β-catenin pathway repressed *FoxD3* expression, which is necessary for Schwann cell development, through *Mitf-M* activation. In conclusion, β-catenin overexpression promotes SCP cell-fate decisions towards the melanocyte lineage.

**Summary statement:** Activation of β-catenin in bipotent Schwann-cell precursors during a specific developmental window, induces MITF and represses FoxD3 to promote melanoblast cell fate at the expense of Schwann cells in limbs.

## INTRODUCTION

Multipotent neural-crest cells (NCC) in vertebrates constitute a transient population of cells arising from the dorsal part of the neural tube (Le Douarin and Kalcheim, 1999) that gives rise to numerous derivatives, such as neuronal and glial cells of the peripheral nervous system (PNS), smooth muscle cells, and melanocytes. Melanocytes produce melanin, a tyrosine-based polymer, in specialized organelles, the melanosomes. Classical melanocytes are pigmented cells, which (i) are found in the skin (dermis or epidermis), (ii) are involved in skin pigmentation, and (iii) are differentiated from melanoblasts derived from late-migrating NCC that have followed the dorso-lateral migratory pathway between the dermamyotome and the overlying ectoderm. These melanoblasts, referred to as first-wave melanoblasts, are specified as early as E8.5, before they start migrating along the dorso-lateral pathway from E10.5 (Petit and Larue, 2016). Between E11.5 and E13.5, most melanoblasts enter the epidermis, where they actively proliferate (Luciani et al., 2011). Between E15.5 and E17.5, epidermal melanoblasts migrate towards the forming hair follicles. In the furry parts of adult mice, most melanocytes are found in the hair matrix, whereas only few interfollicular melanocytes remain in the epidermis after birth (Hirobe, 1984). Epidermal melanocytes are abundant in the hairless parts of the body, such as the tail and paws (Silvers, 1979), except in the palms and soles, which have very few (Kunisada et al., 1998) and (Fig. S1). Melanocytes are considered to be non-classical if they are found in organs other than skin, not involved in skin pigmentation, and/or have not followed the dorso-lateral migratory pathway during development (Colombo et al., 2011). However, two types of non-classical melanocytes involved in skin pigmentation have been found, although they did not follow the dorso-lateral migratory route. One corresponds to a population of cells originating around the time of gastrulation, most likely within the mesoderm, and ultimately residing within the dermis (Kinsler and Larue, 2018). These melanoblasts are referred to as “mesodermal-wave melanoblasts”. The other is derived from Schwann cell precursors (SCPs) and is referred to as second-wave melanoblasts. SCPs are multipotent embryonic progenitors covering all developing peripheral nerves and originate from early ventrally-migrating NCC (Furlan and Adameyko, 2018). Previous studies have shown that a significant number of melanocytes in the skin of the trunk and limbs are produced from SCPs adjacent to spinal nerves that innervate the skin during development. Additionally, it was shown that the glial *versus* melanocyte fate is highly dependent on nerve contact (Adameyko et al., 2009). The authors showed that SCP-derived melanoblasts migrating ventrally from the DRG are specified around E11 in the mouse. While multiple elegant experiments had shown that the melanocytes and Schwann-cells share a common glial-melanogenic bipotent precursor and can be transdifferentiated into each other *in vitro* (Dupin et al., 2000; Dupin et al., 2003; Nitzan et al., 2013b; Real et al., 2006), the factors controlling the cell fate decisions between these two lineages remained unclear. More recent experiments started elucidating the molecular pathways involved in the glial-melanocyte switch. Those bipotent progenitors express various proteins including Sox2, Sox9, Sox10, Fabp, Mitf, Pax3 and FoxD3 (Adameyko and Lallemend, 2010). It has been shown that FoxD3 represses the expression of *Mitf* in zebrafish (Curran et al., 2009), in melanoma cell lines and cultured quail neural crest (Abel et al., 2013; Thomas and Erickson, 2009). Moreover, the downregulation of *FoxD3* is necessary for SCPs to follow a melanocyte fate (Adameyko et al., 2012; Jacob, 2015; Nitzan et al., 2013b).

β-catenin plays critical roles in multiple developmental processes, such as proliferation and cell fate decisions, owing to its dual function in cadherin-dependent cell-cell interactions and as a central component of the canonical Wnt signaling pathway (Aktary et al., 2016; Steinhart and Angers, 2018). Gain-of-function studies have shown induction of cellular proliferation of a number of cell types in transgenic mice expressing stabilized β-catenin (Gat et al., 1998; Imbert et al., 2001; Romagnolo et al., 1999). This pathway influences early melanoblast development, mainly through various common β-catenin/LEF targets, including Myc and Ccdn1, and a major downstream target of β-catenin in the melanocyte lineage, the Mitf-M transcription factor (Luciani et al., 2011). Mitf-M exerts survival and proliferation functions during the expansion of melanoblasts from the neural crest (Carreira et al., 2006; Hornyak et al., 2001) and regulates melanocyte differentiation by inducing the key enzymes of melanogenesis Tyr, Tyrp1, and Dct (Steingrimsson et al., 2004). The deletion of β-catenin specifically in migrating melanoblasts leads to hypoproliferation due to reduced Mitf-M expression (Luciani et al., 2011). Both the temporal and spatial fine-tuning of β-catenin and Mitf-M levels is required to regulate their various downstream targets and generate the required number of melanoblasts at the correct location during development. Apart from its role in neural crest induction and expansion, the Wnt/β-catenin signaling pathway has been implicated in neural-crest cell fate decisions. Mice deficient for both *Wnt1* and *Wnt3a* exhibit a marked deficiency of Dct-positive neural-crest-derived melanoblasts (Ikeya et al., 1997). β-catenin has also been directly associated with melanoblast cell-fate specification in various species using β-catenin gain- and loss-of-function approaches. In zebrafish, injection of *β-catenin* mRNA into a subpopulation of migrating NCCs induces the formation of pigmented cells (Dorsky et al., 1998). In mice, the conditional ablation of *β-catenin* in premigratory NCCs leads to a loss of melanocytes and sensory neurons (Hari et al., 2002), whereas its activation promotes the formation of the sensory neuronal lineage at the expense of other neural-crest derivatives (Lee et al., 2004). A change in cell-fate specification, rather than a proliferation defect, underlies the loss of melanocytes. Moreover, the expression of a constitutive activated form of β-catenin in bipotent cardiac neural-crest cells, known to produce mainly smooth muscle cells and few melanocytes, promotes the melanocyte fate at the expense of the smooth-muscle fate in the *ductus arteriosus* of embryonic hearts, leading to patent ductus arteriosus, a congenital disease (Yajima et al., 2013). Overall, these results demonstrate the essential role of the Wnt/β-catenin pathway in NCC and melanocyte fate determination.

We investigated the influence of β-catenin on the first- and second-wave of melanocyte development. A mouse genetic approach was used to express a conditional mutant of β-catenin (βcatΔex3), known to be hyperactive (Harada et al., 1999), at specific times and in specific neural-crest-cell derivatives using either constitutive or inducible Cre lines under the control of the Tyrosinase promoter (Delmas et al., 2003; Yajima et al., 2006). We observed that constitutive activation of β-catenin led to hyperpigmentation of the paws due to promotion of the melanocyte fate at the expense of the glial fate at the time of SCP specification. At the molecular level, we show that β-catenin overexpression represses *FoxD3* expression through *Mitf,* thereby allowing SCPs to follow a melanocyte fate.

## RESULTS

### Constitutively active β-catenin (βcatΔex3) induces hyperpigmentation of the paws

On a C57BL/6 background, we generated mice producing a constitutively active form of β-catenin (Tyr::Cre/°; βcatex3^flox/+^ = βcatΔex3) in cells of the Tyr::Cre lineage by crossing Tyr::CreA mice (Delmas et al., 2003) with mice harboring a floxed exon 3 of β-catenin (Harada et al., 1999; Yajima et al., 2013). βcatΔex3 mutant mice displayed strong hyperpigmentation of the palms and soles with full penetrance (Figs. 1A, S2A). However, we did not observe strong hyperpigmentation on the back of the paws (Fig. S2A). Palmoplantar hyperpigmentation was already present at birth and was particularly striking at P5 (Fig. 1A). Transversal sections at the metatarsal level of paws from post-natal day 1 (P1) and P5 newborn mice revealed high levels of pigmentation on the ventral side of the βcatΔex3 mutant paws, whereas it was absent from the wild-type (WT) paws (Figs. 1B, S2B). Moreover, this pigmentation was localized in the dermis, directly under the epidermis, as well as more deeply in the palmoplantar mesenchyme.

**Figure 1.**
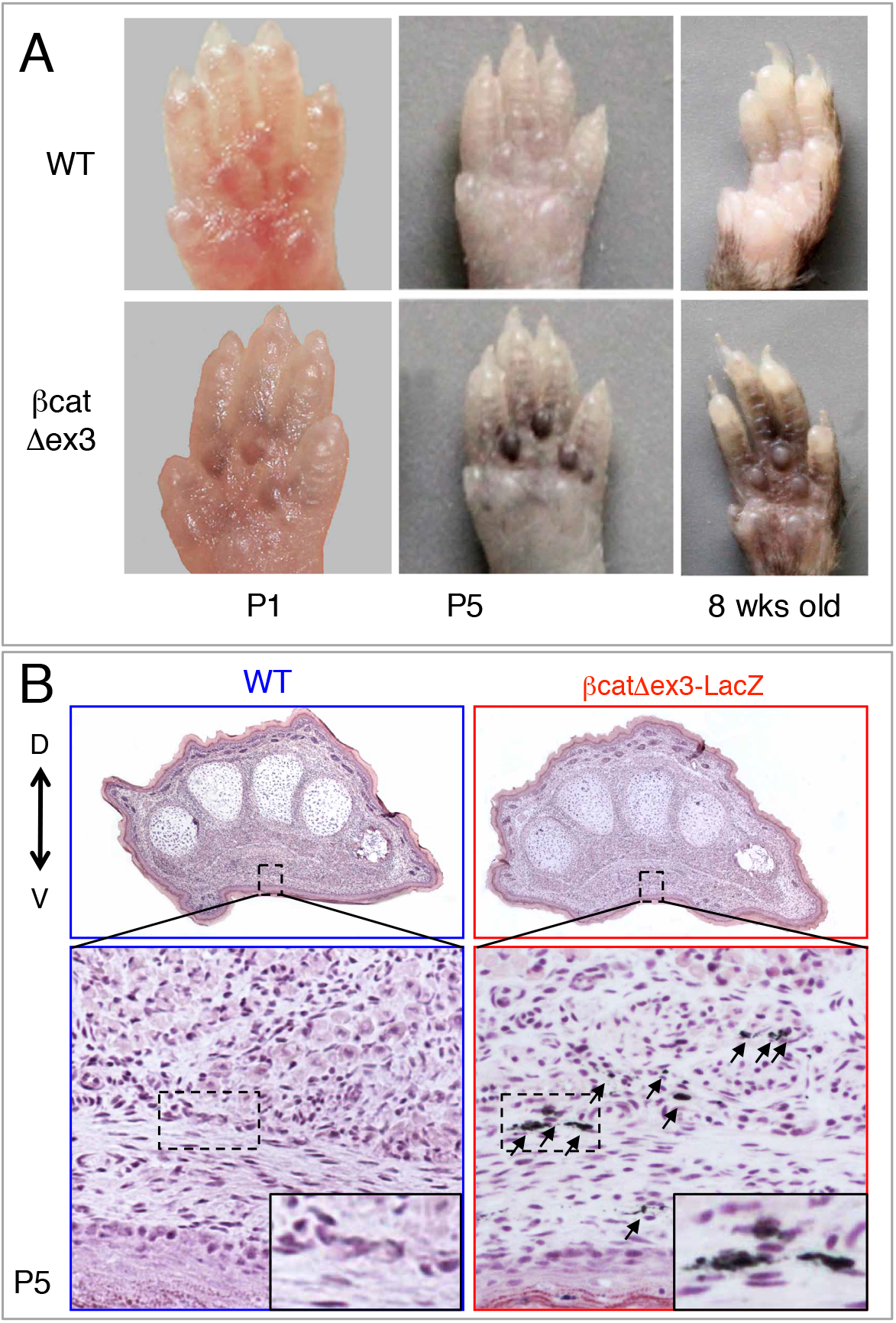
Tyr::Cre/°; β catex3^flox/+^ mice present palmoplantar hyperpigmentation. A) Ventral views of WT and βcatex3 anterior mouse paws in newborns (P1 and P5) and adults. B) Hematoxylin and eosin staining of P5 transversal paw sections. D = dorsal. V = ventral. Arrows point to pigmented cells. WT = (°/°; βcatex3^flox/+^) or (Tyr::Cre; βcatex3^+/+^); βcatΔex3 = (Tyr::Cre/°; βcatex3^flox/+^).

### β-catenin is properly defloxed and activated in βcatΔex3 melanoblasts and melanocytes

The transcriptional activity of βcatΔex3 was previously assessed with the “TOP and FOP” flash luciferase reporter assay and was shown to be five times higher than that of WT β-catenin (Yajima et al., 2013). Deletion of exon3 in βcatΔex3 mice was verified by PCR on genomic DNA extracted from mouse tails containing melanocytes (Fig. S3A,B). We verified the presence of β-catenin in the nucleus, a marker of its stabilization/activation, by immunofluorescence of skin sections during development and after the birth of Tyr::Cre/°; Dct::LacZ (WT-LacZ) and Tyr::Cre/°; βcatex3^flox/+^; Dct::LacZ (βcatΔex3-LacZ) mice using β-galactosidase expression as a melanoblast/melanocyte marker (MacKenzie et al., 1997; Yajima et al., 2013). β-catenin was present in the nucleus of βcatΔex3 melanoblasts in the epidermis of E14.5 embryos whereas it was localized at the membrane in WT mice (Fig. S3C). These results show that β-catenin was properly defloxed and activated in βcatΔex3 melanoblasts and melanocytes.

### The βcatΔex3 mutation does not affect coat color nor truncal melanoblast proliferation

βcatΔex3 mutant mice have no distinctive coat color, ear, or tail phenotype (Fig. S4A). The mutation of β-catenin is induced around E9.0, as the Tyr::Cre transgene begins to be expressed, after dorso-laterally-migrating melanoblasts have been determined. We evaluated the number of WT-LacZ and βcatΔex3-LacZ melanoblasts in the truncal region of E13.5 to E18.5 embryos. From E13.5 to E15.5, the number of melanoblasts was determined on whole mount embryos stained with X-gal in a region localized between the fore- and hindlimbs (ranging from approximately somite 13 to somite 25). There was no significant difference in melanoblast numbers at these stages between WT and mutant embryos (Fig. S4B). At E16.5 and E18.5, truncal melanoblasts were counted on embryo sections immunostained for β-galactosidase (Fig. S4C). Few or no melanoblasts were present in the dermis at these stages, as previously described for WT embryos (Luciani et al., 2011). The presented figures correspond to epidermal and hair-follicle melanoblasts. At E16.5, hair follicles have just initiated invagination from the epidermis while at E18.5, they extend into the dermis and numerous melanoblasts can be found entering and within the hair follicles. There was no difference in melanoblast numbers between WT and mutant mice at these two stages (Fig. S4C). We also investigated melanoblast proliferation in the skin of the trunk using bromodeoxyuridine (BrdU) incorporation assays on embryos collected at E16.5 and E18.5. There was no significant difference in the percentage of BrdU-positive melanoblasts at these stages (Fig. S4D). Overall, these results show that hyperactivation of β-catenin does not influence the development of already determined dorso-laterally-migrating melanoblasts.

### Hyperpigmentation of βcatΔex3 ventral paws is due to an elevated number of melanocytes

X-gal staining of transversal sections of P1 WT-LacZ and βcatΔex3-LacZ paws revealed numerous Dct-positive cells colocalized with strong pigmentation in the mutant palms and soles, whereas they were absent in WT littermates (Fig. 2A,B). X-gal staining also labeled the nerves in the posterior paws (Fig. 2B), but not in the anterior paws (Fig. 2A) (MacKenzie et al., 1997). These nerve-associated Dct::LacZ positive-cells were most likely melanoblasts and/or bipotent SCP and not nerve projections, as we observed a similar pattern of Dct-positive cells colocalized with pigment in both the anterior and posterior paws. The pigmentation pattern in mutant paws was located around the nerves, most likely following nerve projections (Fig. 2B). In the phalanges, pigmentation was strikingly localized around the bones of the digits (Fig. 2B), whereas at the metacarpal/metatarsal level, it was mostly localized under the epidermis (Fig. 2A). These results suggest that ectopic melanocytes are present in the mutant paws.

**Figure 2.**
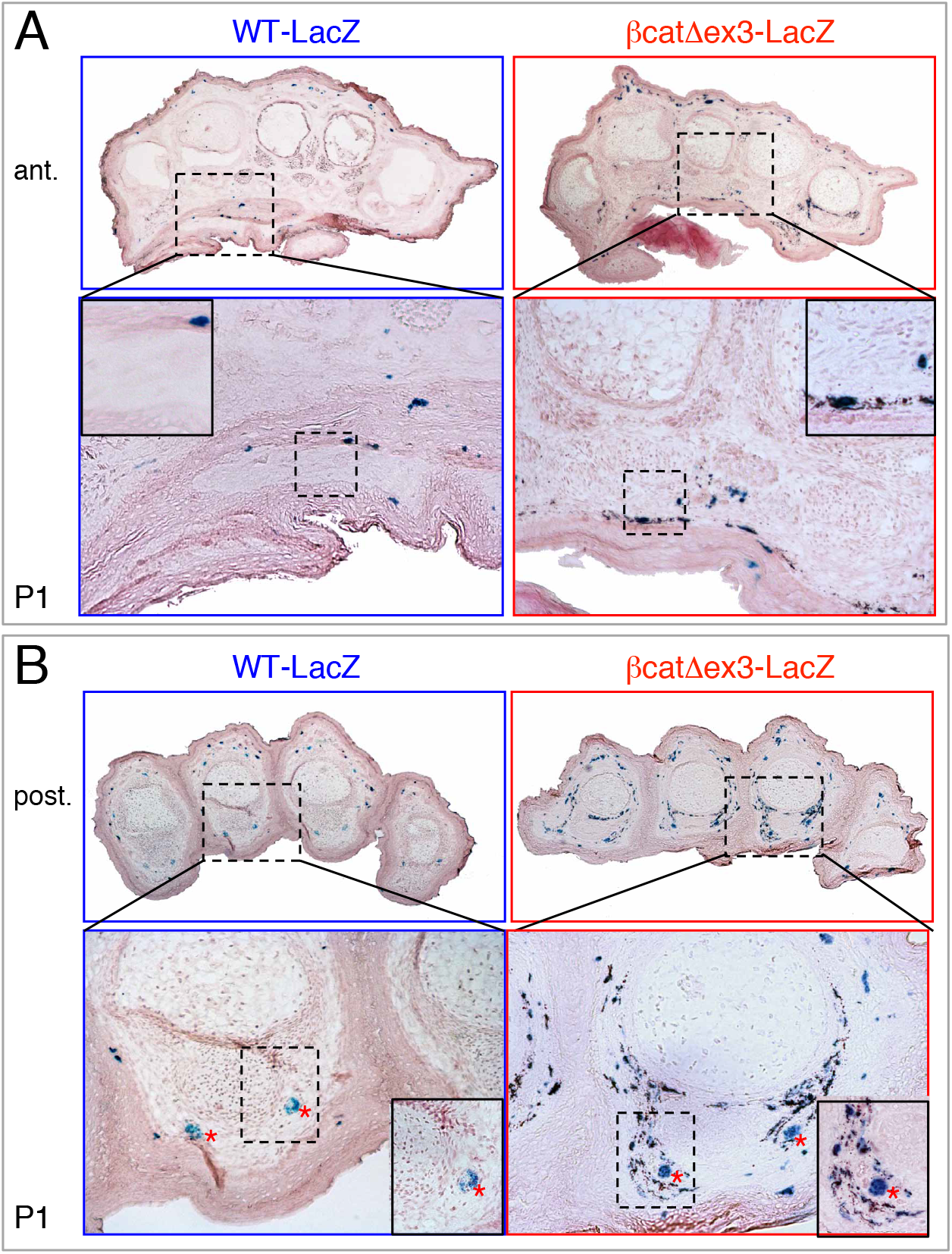
Overexpression of an active form of β-catenin induces hyperpigmented Dct-positive cells on the ventral side of the paws. WT-LacZ and βcatΔex3-LacZ P1 paws were transversally sectioned, and X-gal and eosin stained. (A) Anterior (ant.) and (B) posterior (post.) paws at the metacarpal and phalangeal levels, respectively. In the dermis, mutant paws display high numbers of Dct-positive cells (melanocytes stained in blue, directly under the dermo-epidermal junction of the palmoplantar side of the paws (A) and around the bones of the digits (B). Note that these cells show high accumulation of melanin. In the dermis, WT paws contain a very low number of Dct-positive cells or pigmentation. Note that some nerves are stained in blue in the posterior paws (red asterisk) in WT and mutant paws. WT-LacZ = (°/°; βcatex3^flox/+^; Dct::LacZ/°); βcatΔex3-LacZ = (Tyr::Cre/°; βcatex3^flox/+^; Dct::LacZ/°).

### Hyperpigmentation of βcatΔex3 ventral paws is due to abnormal invasion of melanoblasts during development

The βcatΔex3 paw phenotype was already visible at birth, when the mice are normally unpigmented. It is thus likely the consequence of altered developmental processes. We analyzed the location and number of melanoblasts in E13.5 limbs and paws, when melanoblasts have started their migration to the limbs but have not yet reached the paws. There was no difference in melanoblast distribution between WT and mutant embryos at this stage (Fig. S5A). A difference started to appear at E14.5. Anterior mutant paws displayed melanoblasts ventrally in the palms, as well as few melanoblasts dorsally, whereas they were not present or in very low numbers in WT embryos (Figs. 3A, S5B). There was a statistically significant increase in the number of Dct-positive melanoblasts in the distal region of the ventral limbs, but not in the proximal region of the limb (Fig. 3C). While there was a tendency to increased numbers also on the dorsal side of mutant paws, the difference was not statistically significant (Fig. 3C). No phenotype was yet visible in the posterior paws at this stage (not shown). At E15.5, the phenotype was clearly visible ventrally in mutant paws. Large numbers of melanoblasts were found in the palms and soles and proximal part of the digits, whereas they were mostly absent from the WT paws. Melanoblasts could also be seen in the digits on the dorsal side of the paws (Figs. 3B, S5C). In WT mice, a clear front of migration of melanoblasts was apparent at the junction between the limb and paw (Figs. 3A, S5, black dotted lines). In mutant mice, however, melanoblasts appeared to cross this junction and continue their migration into the palms, soles, and digits. Altogether, these results suggest that constitutively active β-catenin during the establishment of the melanocyte lineage induces melanoblast colonization into the palms and soles.

**Figure 3.**
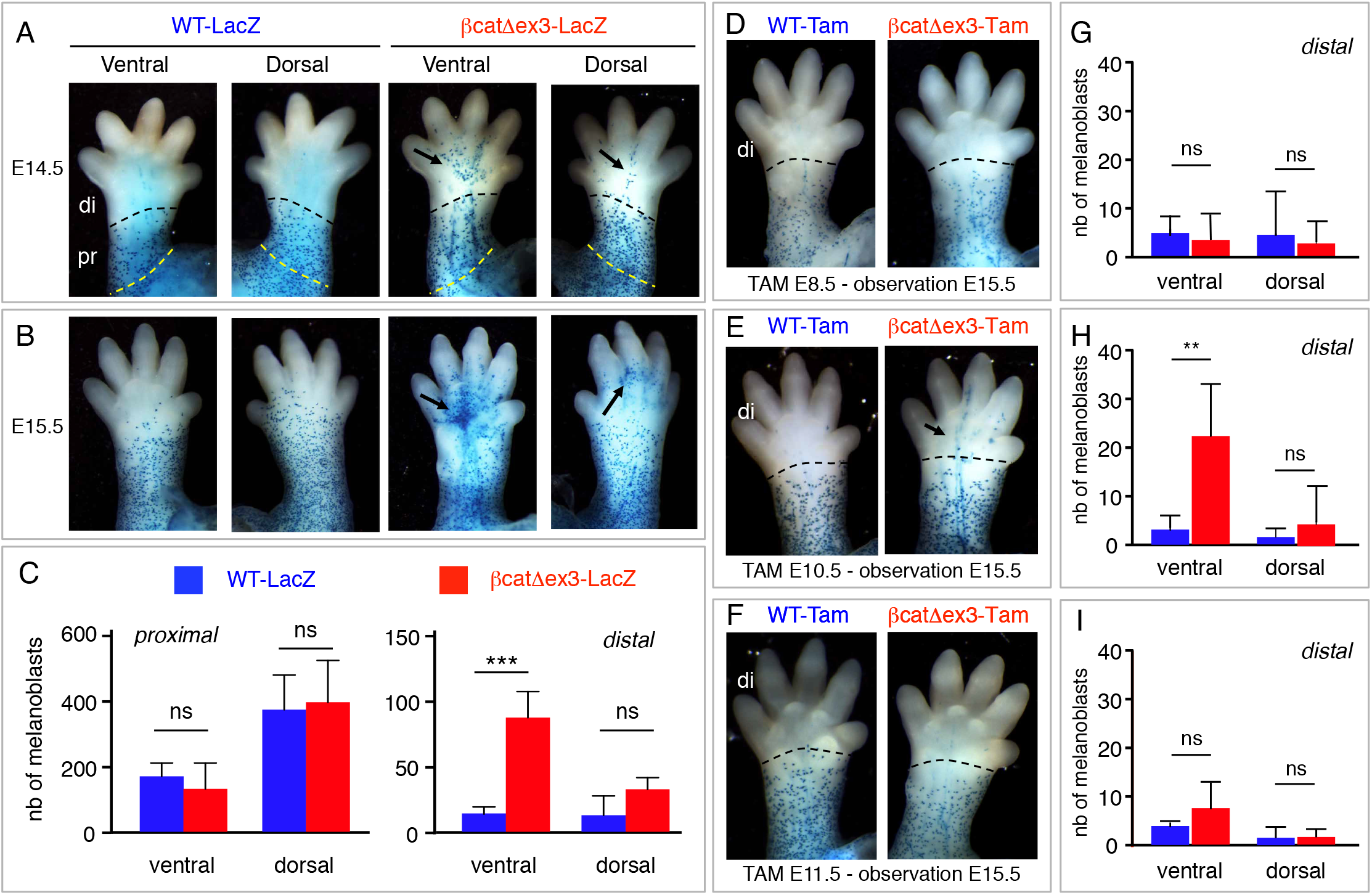
β-catenin favors the specification of SCPs towards melanoblasts. (A-C) The number of melanoblasts is higher on the ventral side of the distal limbs of βcatΔex3 than WT mice. WT-LacZ and βcatΔex3-LacZ E14.5 (A) and E15.5 (B) paws were X-gal stained. Dorsal and ventral views are shown. The number of melanoblasts were estimated at E14.5 (C) in the distal (di) and proximal (pr) region of the limbs that are limited by dashed lines in (A). Arrows highlight ectopic melanoblasts. WT-LacZ = (°/°; βcatex3^flox/+^; Dct::LacZ/°); βcatΔex3-LacZ = (Tyr::Cre/°; βcatex3^flox/+^; Dct::LacZ/°). (D-I) Melanoblast numbers in the paws are increased when β-catenin is activated at E10.5. Ventral views of WT-Tam and βcatΔex3-Tam E15.5 paws induced with 4OH-tamoxifen at E8.5 (D), E10.5 (E), and E11.5 (F) and X-gal stained. The number of melanoblasts was estimated at E15.5 in the distal (di) part of the paw that is delimited by the dashed lines in (D-F) after 4OH-tamoxifen induction at E8.5 (G), E10.5 (H), and E11.5 (I). Arrow in (E) highlights ectopic melanoblasts. No X-gal positive cells were observed at E15.5 when TAM induction was performed at E12.5. WT-Tam = (°/°; βcatex3^flox/+^; Dct::LacZ/°); βcatΔex3-Tam = (Tyr::Cre-ER^T2^/°; βcatex3^flox/+^; Dct::LacZ/°). Using an impaired t-test *** = p-value < 0.001, ** = p-value < 0.01 ns = non significant.

### Melanocytes from the palms and soles originate from the second wave of melanoblasts

Melanocytes are specified from the neural crest around E8.5-E9.0 (Le Douarin and Kalcheim, 1999), while they seem to specify from SCPs around E10.5-E11.5 (Fig. S6) (Adameyko et al., 2009; Van Raamsdonk and Deo, 2013). We used temporal induction of βcatΔex3 to reveal the origin of the melanoblasts invading the soles and palms and leading to the hyperpigmentation phenotype. We generated Tyr::CreER^T2^/°; βcatex3^flox/+^ mice (βcatΔex3-Tam), induced activated β-catenin at either E8.5, E10.5 or E11.5 with 40H-tamoxifen, and evaluated the location and number of melanoblasts in the distal part of the limbs at E15.5. The tamoxifen induction at E8.5 and E11.5 appeared to affect neither the number nor localization of melanoblasts at E15.5 in the distal region of the ventral paws (Fig. 3D,F,G,I). However, tamoxifen induction at E10.5 resulted in a clear increase in the number of melanoblasts in the distal region of the ventral paws (Fig. 3E,H). These results suggested that SCPs actually specify into melanocytes as early as E10.5 and that hyperactivation of β-catenin in those bipotent progenitors at that specific time promotes the melanoblast fate. We thus estimated the number of Schwann cells (Gfap-positive cells) and melanoblasts (Mitf-positive cells) in βcatΔex3 limbs. Expression of βcatΔex3 led to an increased number of melanoblasts and a decreased number of Schwann cells in the palms (Fig. 4). These results suggested that the expression of a constitutively active form of β-catenin in glial-melanogenic bipotent progenitors at the time of their fate determination promoted their differentiation into melanoblasts of the second wave at the expense of glial cells. Because SCPs are located along and migrate with axons of peripheral nerves, the ectopic melanocytes observed in the paws of βcatΔex3 mice would likely have migrated away from these nerves.

**Figure 4.**
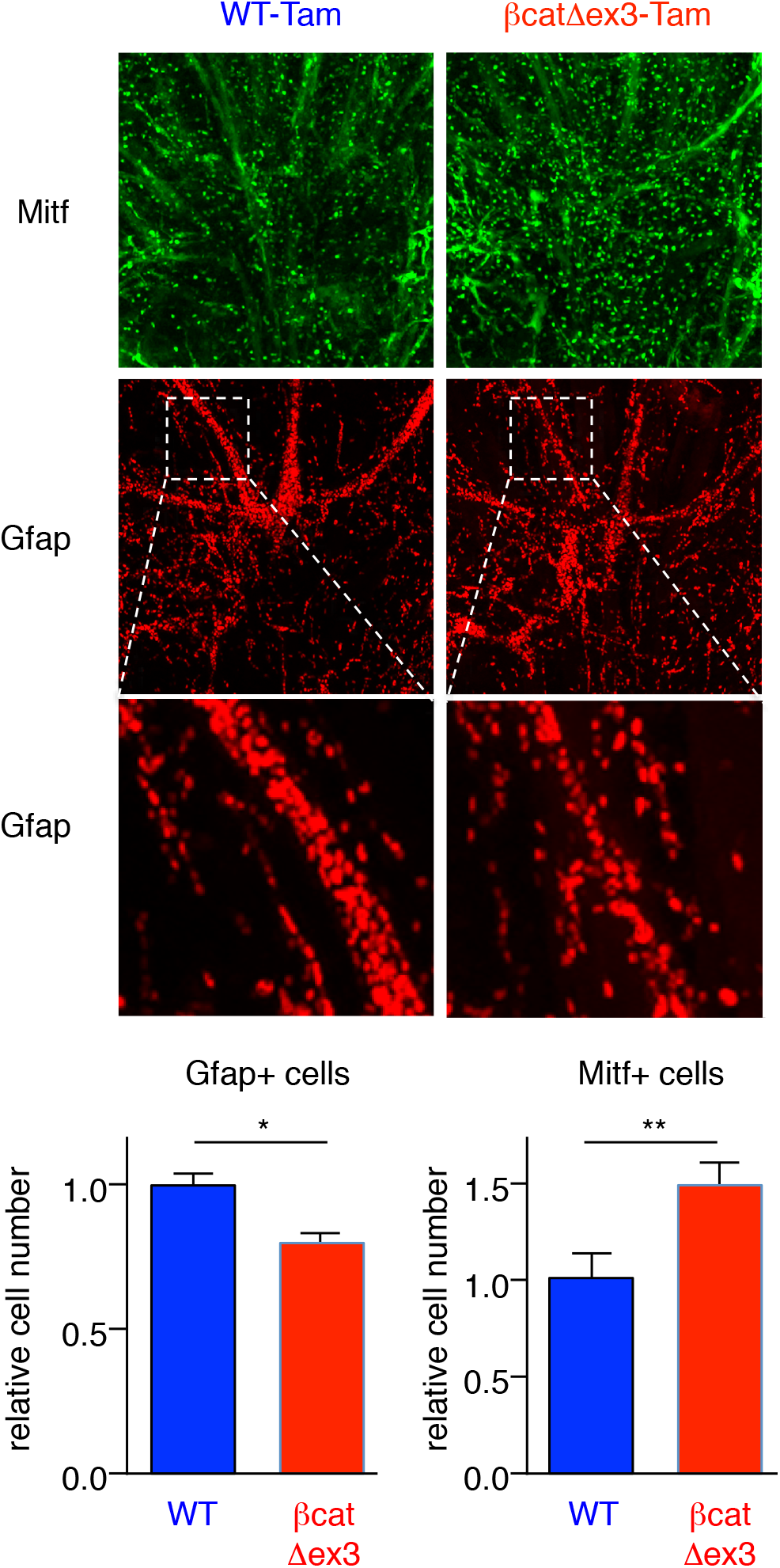
The number of paw melanoblasts increases at the expense of glial cells when β-catenin is activated at E10.5. Ventral views of WT-Tam and βcatΔex3-Tam E15.5 anterior paws from embryos induced with 4OH-tamoxifen at E10.5. Immunostainings for Mitf-M (green) and Gfap (red). A zoom at the level of the nerve is presented and highlights the reduction of Gfap positive cells in βcatΔex3 paws compared to WT. WT-Tam = (°/°; βcatex3^flox/+^); βcatΔex3-Tam = (Tyr::Cre-ER^T2^/°; βcatex3^flox/+^). The relative amounts of Gfap positive (Gfap +) and Mitf positive (Mitf +) cells are shown (WT *vs.* βcatΔex3). Statistical analysis was performed using an unpaired t-test. Error bars correspond to SEM. *p < 0.05 and **p<0.01.

### β-catenin promotes the SCP-derived melanocyte fate through Mitf repression of FoxD3

The downregulation of *FoxD3* in SCPs is necessary to allow emergence of melanocyte cells (Adameyko et al., 2012; Jacob, 2015; Nitzan et al., 2013a) prompting us to ask whether activation of β-catenin signaling affected *FoxD3* expression. Constitutive activation of β-catenin signaling by knocking down *APC* using an siRNA in HEI-193 human schwannoma cells resulted in a significant decrease of *FOXD3* mRNA level compared to control scrambled siRNA (siScr) transfected cells (Fig. 5A). As a control, we showed that under the same conditions the levels of *AXIN2* mRNA, a well-known downstream target of β-catenin, was induced (Fig. 5B). It has previously been shown that *FOXD3* overexpression in melanoma cell lines or cultured quail neural crest cells resulted in repression of *MITF* expression (Abel et al., 2013; Thomas and Erickson, 2009). In a converse experiment, we show here that siRNA-mediated *MITF* silencing in 501mel and SK28 human melanoma cells led to upregulation of *FOXD3* (Fig. 5C-F). ChIP-seq in 501mel cells revealed that MITF occupied several sites at the *FOXD3* locus in a putative distal enhancer. One of these sites was co-occupied by SOX10 and marked by H3K27ac, BRG1 and H2AZ (Fig. 5G). MITF ChIP-seq in primary human melanocytes also showed MITF binding to the distal enhancer in particular at the site co-occupied by SOX10, but in addition, binding to a site in the proximal *FOXD3* promoter (Webster et al., 2014). At each site, a consensus E-box sequence was present along with a SOX10-binding motif at the distal enhancer. Moreover, these binding sequences were present at the otherwise well-conserved syntenic regions at the mouse *Foxd3* locus (Fig. 5G). Taken together, these observations strongly suggested the presence of a reciprocal regulatory feedback loop in the melanocytic lineage where FOXD3 repress *MITF* and MITF repress *FOXD3*. Since the level of MITF expression and activity depends on numerous factors in the melanocytic lineage, this equilibrium may be rapidly shifted in favor of MITF when one of these molecular pathways, such as Wnt/β-catenin, is induced leading to decreased FOXD3 levels and altered cell fate (see schematic on Fig. 6).

**Figure 5.**
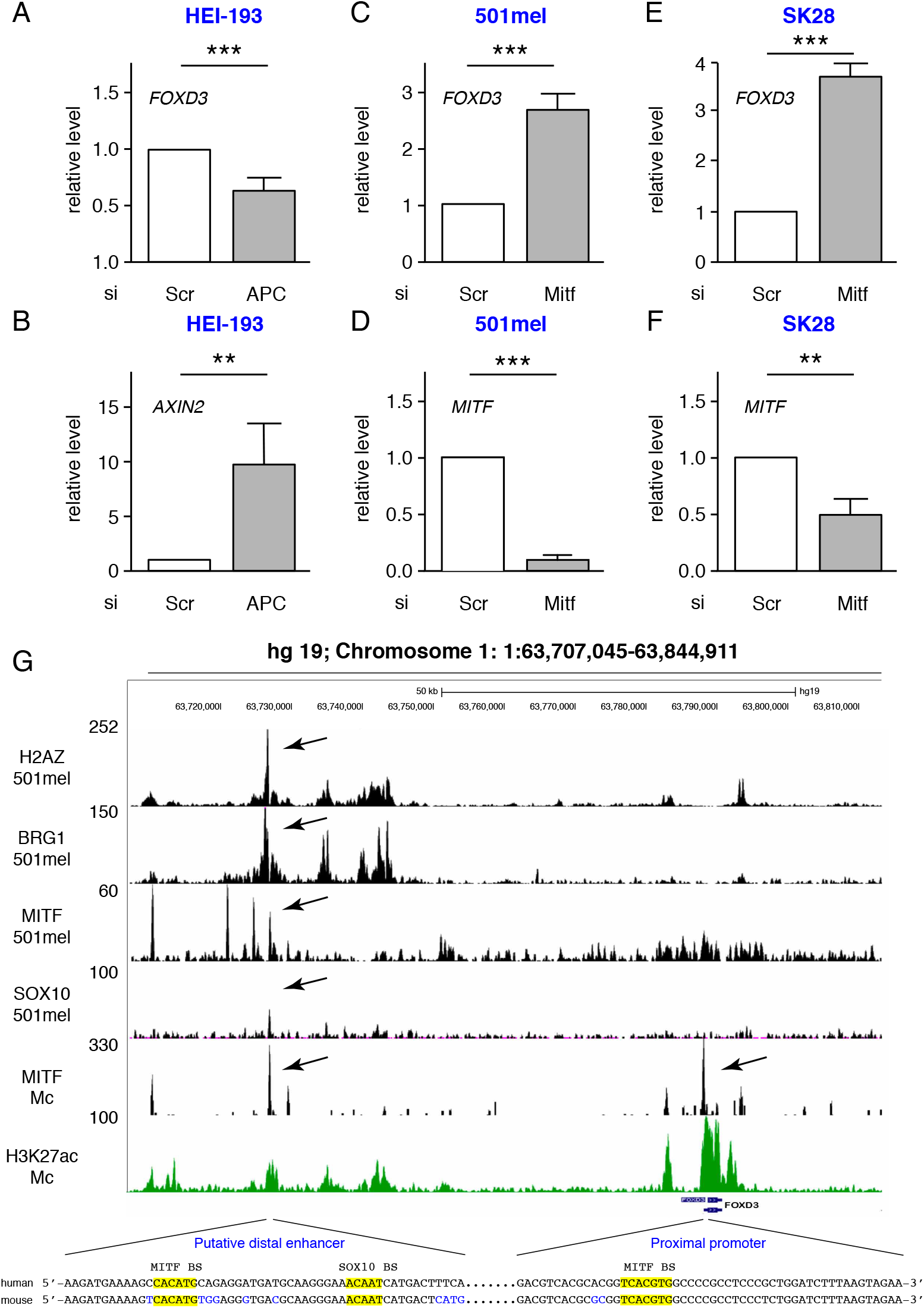
MITF represses *FOXD3* expression. (A,B) The relative amounts of *FOXD3* and *AXIN2* were determined by RT-qPCR from the HEI-193 schwannoma cell line after siRNA mediated knockdown of APC. (C-F) The relative amounts of *FOXD3* (C,E) and *MITF* (D,F) were determined by RT-qPCR in 501mel and SK28 human melanoma cell lines after siRNA mediated knockdown of *MITF,* respectively. (G) UCSC screenshot of ChIP-seq data at the FOXD3 locus. Shown are ChIP-seq data for H2AZ, BRG1 MITF and SOX10 in 501mel melanoma cells as previously described (GSE GSE61967) and for H3K27ac from GSM958157 (Laurette et al., 2015). MITF ChIP-seq in primary melanocytes (GSE50686) is from (Webster et al., 2014). Binding sites are indicated by arrows in the proximal promoter in primary melanocytes (Mc) and in a putative distal enhancer in Mc and 501mel cells. The DNA sequences under the peaks are shown along with the syntenic regions from mouse. MITF and SOX10 binding sites (BS) are highlighted in yellow. Each of these BS are bound by BRG1 and H2AZ; additional marks of regulatory sequences. Statistical analysis was performed using the unpaired t-test. Error bars correspond to SD. **p < 0.01 and ***p<0.001.

**Figure 6.**
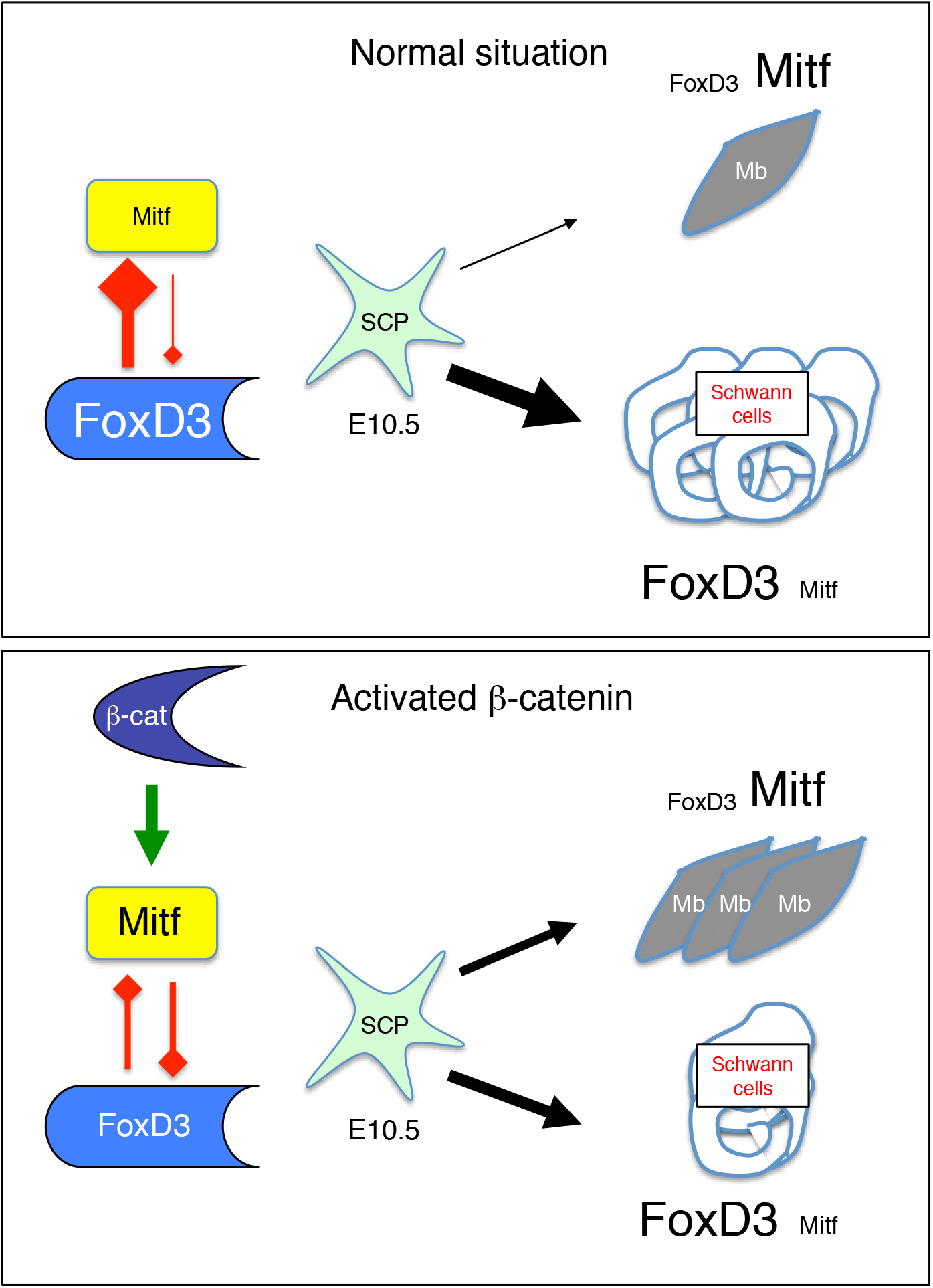
Schematic of the determination of SCP to generate Schwann cells and melanoblasts.

## DISCUSSION

Here, we show that a constitutively active form of β-catenin (βcatΔex3) differentially affects melanoblast development in the trunk and paws. In the trunk region, expression of βcatΔex3 did not induce any major defects in developing melanoblasts, whereas it induced strong palmo-plantar hyperpigmentation of the paws. This hyperpigmentation was due to the abnormal presence of melanocytes derived from the second wave of melanoblasts. Melanoblasts migrating in the palms and soles of the mutant mice were seen as early as E14.5, whereas they were mostly absent in WT mice. These results show that once specified, βcatΔex3 does not influence the development of melanoblast of the first wave, but instead controls SCP cell-fate decisions between glial and melanocyte lineages in the ventral migratory pathway. According to these results, the contribution of SCPs to melanocytes in the adult appeared to be restricted to the limbs.

### Hyperpigmentation of the paws

Hyperpigmentation of the palms and soles was already described in humans and mice after cell non-autonomous induction. Human palmoplantar fibroblasts express the Wnt/β-catenin signaling inhibitor DKK1, which inhibits melanocyte function and growth by regulating β-catenin (Yamaguchi et al., 2004; Yamaguchi et al., 2008). Downregulation of β-catenin leads to the inhibition of Mitf-M expression, and of its downstream target Tyrosinase, the key enzyme of melanogenesis. Increasing β-catenin levels in SCP-derived melanocytes may counteract the effects of Dkk1 in palmoplantar skin, promoting melanocyte differentiation after inducing Mitf and Tyrosinase. Overexpression of Kitl (Steel factor) in the basal layer of the epidermis in mice induces palmoplantar hyperpigmentation (Kunisada et al., 1998). The authors found melanoblasts in the footpads of E16.5 mutant embryos, whereas they were not present in WT littermates. As Kit signaling is involved in melanoblast migration, they proposed that increased Kit signals promote migration of melanoblasts throughout the entire paw epithelium. This explanation is certainly valid. However, Kit signaling in melanocytes indirectly regulates β-catenin, through the PI3K pathway, and Mitf-M, through the MAPK pathway. Thus, in keeping with the results obtained here, an alternative and/or complementary explanation for the palmoplantar hyperpigmentation is enhanced melanoblast specification from SCPs.

### Specification

As previously mentioned, β-catenin is involved in cell-fate specification, a process involving complex combinations of cell intrinsic and extracellular signals that need to be correctly delivered in time and space. The role of β-catenin in the specification of first wave melanocytes has been clearly demonstrated. The inactivation of β-catenin in NCC prior to melanoblast specification using Wnt1::Cre shows that β-catenin is essential for the generation of melanoblasts. The absence of β-catenin apparently does not impair early SCP specification, as specific markers are produced (Hari et al., 2002). Thus, SCPs and second wave melanocytes still form in these animals. This series of experiments showed the critical function of Wnt signaling in driving early melanoblast specification and could explain the absence of first-wave melanocytes (*i.e.* migrating dorso-laterally), but the importance of β-catenin in the generation of second wave melanocytes was still unknown. As Schwann cells and second wave melanocytes share a common SCP precursor, we hypothesized that β-catenin in the βcatΔex3 mutant mice is activated in SCPs that migrate via the ventral pathway, altering their fate and promoting their differentiation into melanocytes. Whereas neural progenitors and glial cells express the Foxd3 transcription factor, it is not expressed in melanoblasts (Kos et al., 2001). As Mitf is the key transcription factor specifying the melanocyte lineage and knowing that SCPs express Foxd3, Mitf and Sox10, it is likely that SCP fate is governed by the relative amounts/activities of Foxd3 and Mitf. In agreement with this hypothesis, constitutive activation of β-catenin in Schwannoma cells led to FOXD3 repression, whereas *MITF* silencing up-regulated FOXD3 expression in melanoma cell lines. Moreover, MITF binds to regulatory elements at the *FOXD3* locus in human melanoma cells and primary melanocytes and may therefore directly inhibit its expression. In contrast, overexpression of FOXD3 in melanoma cell lines represses *MITF* expression (Abel et al., 2013; Thomas and Erickson, 2009). Together these observations support the idea that a direct and reciprocal negative regulation of FOXD3 and MITF expression can affect SCP fate. This model is reminiscent of the reciprocal negative regulation seen with MITF and JUN that affects the phenotype switch between melanocytic and undifferentiated melanoma cell states (Riesenberg et al., 2015). Based on these observations, we propose that high β-catenin levels in SCP at the time of their specification increases *Mitf* expression, hence repressing *FoxD3* expression and enhancing melanocyte specification at the expense of glia. Such a model is supported by the reduced numbers of Gfap-positive cells and increased numbers of Mitf-positive or Dct-positive cells observed in the paws of βcatΔex3 mice, suggesting that a cell fate switch occurred.

### Acral melanoma

Although the number of melanocytes in the soles of the feet and palms of the hands are very limited, these cells may transform in acral melanoma (ALM). ALM and nodular melanoma (NM) are more aggressive than superficial spreading melanoma (SSM). The percentage of ALM is higher in Asians (50%) than in Caucasians (10%). This is because NM and SSM are very rare in Asians, but the risk to develop an ALM appears to be similar between Asians and Caucasians. At the molecular level, the main mutations in ALM and non-ALM are similar; they include mutations in the *BRAF, NRAS, NF1,* and *KIT* genes, but the proportions are different (Moon et al., 2018; Zebary et al., 2013). NM and SSM arise from melanocytes determined from the first wave of melanoblasts, while ALM arises from melanocytes derived from the second wave of melanoblasts. While sun exposure is a well-established cause for melanoma development, the soles and palms are non-sun-exposed regions, raising the issue of the importance of the embryonic origin of melanocytes in melanomagenesis and how this may influence their aggressivity when transformed.

### Conclusion

β-catenin appears to play a complex role in the melanocyte lineage, depending on tight regulation of its levels and time and place of induction. We show here that expression of βcatΔex3 after specification of the melanoblasts of the first wave in Tyr::Cre-and Tyr::CreER^T2^-expressing cells does not appear to affect melanoblast development in the dorso-lateral pathway, but favors melanoblast specification in the ventral pathway.

## MATERIALS AND METHODS

### Transgenic mouse generation and genotyping

Animal care, use, and experimental procedures were conducted in accordance with recommendations of the European Community (86/609/EEC) and Union (2010/63/UE) and the French National Committee (87/848). Animal care and use were approved by the ethics committee of the Curie Institute in compliance with the institutional guidelines.

Mice with conditional constitutive stabilization of β-catenin were generated by mating Tyr::CreA and Tyr::Cre-ER^T2-Lar^ (designated in the text as Tyr::Cre-ER^T2^) transgenic mice (Delmas et al., 2003; Yajima et al., 2006) with animals homozygous for a floxed allele of β-catenin, with LoxP sites flanking exon 3 (Δex3) (Harada et al., 1999). Transgenic mice were maintained on a pure C57BL/6J background (backcrossed at least 10 times). All animals were housed in specific pathogen-free conditions in the animal facility. Mice were genotyped using DNA isolated from tail biopsies using standard PCR conditions. The Tyr::Cre transgene (0.4 kb fragment) was detected by PCR, as previously described (Delmas et al., 2003). For detection of the floxed (570bp) and WT (376bp) alleles of the β-catenin gene, PCR amplification was carried out with the forward primer (LL523) 5’-GAC ACC GCT GCG TGG ACA ATG-3’ and the reverse primer (LL524) 5’-GTG GCT GAC AGC AGC TTT TCT G-3’. The forward primer (LL667) 5’-CGT GGA CAA TGG CTA CTC A-3’ and the reverse primer (LL668) 5’-CTG AGC CCT AGT CAT TGC AT-3’ were used for detection of the WT (715bp) and deleted (450bp) alleles of the β-catenin gene. The PCR conditions were as follows: 5 min at 94°C followed by 35 cycles of 20 s at 94°C, 30 s at 56.5°C, 45 s at 72°C, and a final extension of 10 min at 72°C.

### Tamoxifen injection

Pregnant C57BL/6J mice were injected intraperitonnaly at E8.5, E10.5 or E11.5 with tamoxifen (Sigma) diluted in corn oil. An amount of 0.5mg of tamoxifen was injected for 20g of body weight. This dose of Tamoxifen was not optimal but higher doses induced embryonic death and resorption of the embryos.

### Histology

Mice were crossed with Dct::LacZ (MacKenzie et al., 1997) and the resulting embryos collected at various times during pregnancy. Embryos were stained with X-gal, as previously described (Delmas et al., 2003). Paws of new-born mice at P1 were dissected, washed in PBS, and fixed by incubation in 0.25% glutaraldehyde in PBS for 50 min at 4°C. They were then incubated in 30% sucrose/PBS overnight, followed by 30% sucrose/50% OCT/PBS for 5h and embedded in Optimal Cacodylate Compound (OCT). Cryosections (8μm thick) were stained either with heamatoxylin and eosin or X-gal as follows: they were washed twice in PBS at 4°C, and incubated twice, for 10 min, in permeabilization solution (0.1M phosphate buffer pH7.3, 2mM MgCl_2_, 0.01% sodium deoxycholate, 0.02% NP-40) at room temperature. They were then incubated in staining solution (0.4mg/mL 5-bromo-4-chloro-3-indolyl-D-galactosidase, 2 mM potassium ferricyanide, 2 mM potassium ferrocyanide, 4 mM MgCl_2_, 0.01% sodium deoxycholate, and 0.02% NP-40 in PBS) overnight at 30°C. Sections were post-fixed in 4% PFA overnight at 4°C, washed in PBS, and stained with eosin. Paws of newborn mice at P5 were fixed in 4% PFA, dehydrated, and embedded in paraffin by standard methods. Paraffin sections (7μm thick) were stained with eosin.

### Immunostaining

Mice were crossed with Dct::LacZ and the embryos collected at various stages of development. Newborn skin was dissected from the back of the mice. Embryos and skins were washed in PBS and fixed by overnight incubation in 4% PFA. They were then incubated in 30% sucrose/PBS overnight, followed by 30% sucrose/50% OCT/PBS for 5 h and embedded in OCT. Cryosections (7 μm thick) were washed with PBS-Tween 0.1% (PBT) for 10 min. Antigens were then retrieved by incubation for 20 min in citric acid buffer (pH 7.4) at 90°C. Non-specific binding was blocked by incubation with 2% skimmed milk powder in PBT. Sections were incubated overnight at 4°C with various primary antibodies. Rabbit polyclonal antibody against β-catenin (Abcam ab6302) and chicken polyclonal antibody against β-galactosidase (Abcam ab9361) were used. Sections were washed three times in PBST for 5 min each and incubated with secondary antibodies for 1 h at 37°C. The secondary antibodies used were donkey Alexa 488-anti-rabbit and donkey Alexa 555-anti-chicken (Molecular Probes). Sections were incubated in DAPI for 10 min, washed three times in PBT, for 10 min each, and mounted in mounting media containing N-propylgalate. Conventional fluorescence photomicrographs were obtained with a Leica DM IRB inverted routine microscope.

### Whole mount immunostaining

E13.5 and E15.5 embryos paws were collected and fixed in 4% PFA-PBS pH7.5 (Euromedex) for 6 hours prior washing them three times in PBT. Paws were dehydrated in a series of PBS/methanol incubation (25%, 50%, 75% and 100%) for 10min each. Paws were incubated 24 hours in methanol at 4°C prior bleaching them for 24h in a mixture of 1/3 H_2_O_2_ and 2/3 methanol 30%. Paws were washed three times in methanol prior post-fixed them overnight in 1/5 DMSO and 4/5 pure methanol. Paws were sequentially rehydrated in PBS/methanol (75%, 50%, 25% and 0%) for 10 min each prior washing them twice in PBT.

Paws were incubated overnight at room temperature in PBS containing 5% Donkey serum, 1% BSA and 20% DMSO. After blocking, paws were incubated with primary antibody(ies) in the blocking solution at 1/1,000 for five days at RT. Primary antibodies were rabbit against neurofilament (Abcam ab9034), mouse against Gfap (Sigma C9205 & Cell Signalling Technology 3670) and goat against Mitf (R&D system AF5769). Secondary antibodies were diluted in the blocking solution: alexaFluor 555 (Invitrogen A-31572), alexaFluor488 (Invitrogen A21202) and alexaFluor633 (Invitrogen A-21082) for overnight at RT. Staining was ended after incubation of the paws in Dapi for 4 hours at RT. Embryos were dehydrated in PBS/methanol (25%, 50%, 75% and 100%) for 10min each at RT. Chambers made by 1mm thick Fastwell (Sigma) coated on a glass slide was used to incorporate the paws. Each paw was fixed on to the glass slide with 1% NuSieve Agarose (Sigma) and covered with methanol. After three washes with methanol, paws were incubated twice for 5min with methanol 50% BABB (1/3 benzylalcohol and 2/3 benzylbenzoate, from Sigma), and then three times in pure BABB for 5min each (or until the sample is cleared). The chamber was closed with coverslip and sealed with nail polish prior examination under the microscope.

### Confocal imaging and ImageJ treatment for 3D reconstruction

Z-sections were acquired every 5 μm for the dorsal and ventral part of the limb with a Confocal Leica SP5 microscope. Then, the pluggin PureDenoise was used on the stack to increase the signal, and finally the filter substract background (20) was used to remove the remaining background. 3D reconstructions were performed from stacks containing the same number of sections and the same biological structures in WT and mutants, using 3D project in *ImageJ* without interpolation.

### BrdU labelling

Melanocyte proliferations were analyzed using BrdU labelling *in vivo* on embryos at various stages of development. BrdU (100 μg/mL, BD Biosciences) was injected intra-peritonally into the pregnant mother 2 h before sacrifice, in the form of two 50-μg/mL injections administered at 20-min intervals. Embryos were collected for immunohistochemistry. They were fixed and stained, as described above, with mouse monoclonal anti-BrdU antibody (BD Biosciences) and chicken polyclonal anti-β-galactosidase antibody (Abcam). Donkey Alexa 488-anti-mouse and donkey Alexa 555-anti-chicken (Molecular Probes) were used as secondary antibodies.

### Melanoblast counts on the paws

Pictures of Xgal stained paws were taken using a binocular magnifying glass with a 1x objective. Proximal area (from the body to the migrating front, between the dotted yellow and black lines) and distal area (after the black dotted line) were delimited on the picture. Blue dots (melanoblasts) were counted using *ImageJ* software. At least 5 embryos were counted for each genotype at each stage in both areas.

### Cell culture and siRNA-mediated knockdown

501mel and SK28 human melanoma cell lines were grown in RPMI 1640 media (GIBCO) supplemented with 10% FCS (GIBCO) and 1% Penicillin-Streptomycin (GIBCO). HEI-193 Schwannoma cells were grown in DMEM media (GIBCO) supplemented with 10% FCS (GIBCO) and 1% Penicillin-Streptomycin (GIBCO). Cells were maintained at 37°C in a humidified atmosphere containing 5% CO2. siRNA targeting human MITF (M-008674) and APC (L-003869) were purchased from Dharmacon. Si Scramble (siSCR), with no known human targets, was purchased from Eurofins Genomics. Cells were transfected with 100 pmol siRNA or siScr with Lipofectamine2000 (Invitrogen) and assayed for mRNA expression 48h post-transfection.

### RNA extraction and RT-qPCR

Total RNA was extracted from cell lines using the miRNeasy kit (Qiagen). M-MLV reverse transcriptase (Invitrogen) was used according to the manufacturer’s protocol to synthesize cDNA from 1 μg total RNA in combination with random hexamers. Quantitative RT-PCR was performed with the iTaq universal Sybrgreen Supermix (BIORAD) and primers listed below, using a QuantStudio 5 thermocycler (Applied Biosystem) in a final reaction volume of 25 μL under the following conditions: 95°C for 1.5min, 40 cycles of 95°C for 30s, 60°C for 60s, with a final melting curve analysis. Relative expression was determined by the comparative ΔΔCt method. PCR primers: FOXD3 f: 5’-CAT CCG CCA CAA CCT CTC-3’; FOXD3 r: 5’-CAT ATG AGC GCC GTC TG-3’; MITF f: 5’-CTA TGC TTA CGC TTA ACT CCA-3’; MITF r: 5’-TAC ATC ATC CAT CTG CAT ACA G-3’; AXIN2 f: 5’-CCT AAA GGT CGT GTG TGG CT-3’; AXIN2 r: 5’-GTG CAA AGA CAT AGC CAG AAC C-3’; TBP f: 5’-CAC GAA CCA CGG CAC TGA TT-3’; TBP r: 5’-TTT TCT TGC TGC CAG TCT GGA C-3’.

### Statistical analysis

Statistical tests are detailed in the figure legends. All data are presented as mean ± SEM. Statistical analyses were performed with Prism 5 software (GraphPad).

## ACKNOWLEDGEMENTS

We are grateful to Dominique Lallemand for providing HEI-193 human schwannoma cell line. We thank the teams caring for the imaging, histology and animal colony facilities of the Institut Curie, especially Pauline Dubreuil and Mirella Miranda.

